# ZNF185 expression is negatively regulated by CTCF and promotes endometrial cancer growth

**DOI:** 10.64898/2026.06.22.733662

**Authors:** Siting Yan, Sarah Ho, Rachel Lin, Quinlyn Satava, Cynthia Metierre, Tinuoye Winjobi, Melissa Vellozzi, Mehdi Sharifi Tabar, John EJ Rasko, Charles G Bailey

## Abstract

CCCTC-binding factor (*CTCF*) is frequently mutated in endometrial cancer, resulting in genetic haploinsufficiency that contributes to tumour progression. We previously showed that depletion of *CTCF* disrupted cell polarity in KLE endometrial cancer spheroids; however, the implications for gene dysregulation and endometrial cancer pathophysiology remains poorly understood. ZNF185, an actin-associated and LIM domain-containing protein involved in cytoskeletal remodelling, was identified as a dysregulated target following *CTCF* haploinsufficiency. In this study, shRNA-mediated knockdown of *CTCF* was used to model haploinsufficiency in endometrial cancer cells, leading to the identification of a previously unrecognised isoform of ZNF185, named ZNF185B. Unlike the full-length protein, ZNF185B lacked co-localisation with F-actin and exhibited a diffuse cytoplasmic distribution, and ZNF185B was significantly upregulated in *CTCF*-depleted endometrial cancer cells and in an auxin-inducible degron model in a dose-dependent manner. Functional studies demonstrated that depletion of ZNF185 expression reduced endometrial cancer cell proliferation and clonogenic potential. Together, these findings identify ZNF185B as a novel isoform negatively regulated by CTCF protein dosage and establish ZNF185 as a requirement for endometrial cancer cell proliferation. Our results suggest that dysregulated ZNF185 expression is a crucial downstream consequence of CTCF haploinsufficiency and may contribute to tumour progression in endometrial cancer.

## Introduction

Endometrial cancer (EC) is the most common gynaecological malignancy in developed countries, with a high incidence (∼one in every 2,000 women^1^) and rising mortality. EC develops in the inner lining of the uterus, and is broadly classified into two pathogenetic types, LJ and LJ, which differ in histology, molecular mechanism, clinical behaviour and prognosis.^2^ However, these traditional classifications now have their limitations due to the considerable morphological and molecular heterogeneity in endometrial cancer.^3^ Type LJ EC, which is typically endometrioid in histology, accounts for the majority of cases. It is estrogen-driven and highly associated with obesity, but with a more favourable overall survival rate. In contrast, Type LJ EC is less common, estrogen-independent, and characterised by serous or clear cell carcinoma. The Type LJ EC incidence is rising in older age and it is more aggressive with poorer outcomes.^2^ These clinicopathological distinctions are also reflected by multiple somatic gene mutations. The genetic landscape of endometrial cancer provides the molecular basis for its initiation and progression. Endometrioid carcinoma is characterised by recurrent somatic mutations in *PTEN*, *PIK3CA*, *PIK3R1*, *ARID1A*, *KRAS*, *CTNNB1*, *CTCF*, *RPL22* and *ARID5B*,^4, 5^ whereas serous subtype carcinoma is largely driven by *TP53* mutations, with additional *PIK3CA* and *PPP2R1A* mutations.^5, 6^ Among 127 significantly mutated genes (SMGs) within different tumour types, *CTCF*, encoding CCCTC-binding factor, has been identified as a SMG in endometrial carcinoma.^4^ *CTCF* is mutated in approximately a quarter of all endometrioid carcinomas, leading to genetic haploinsufficiency.^7^ *CTCF* mutations also occur in other cancers including breast cancer, prostate cancer, Wilms’ tumour, and lymphoblastic leukaemia, but at a lower frequency.^8, 9^

CTCF is a highly conserved protein characterised by a central 11-zinc finger (ZF) domain. It is a multifunctional DNA-binding protein, known as the ‘master weaver of the genome’,^10^ playing diverse roles in chromatin architecture organisation, enhancer and promoter insulation, transcriptional regulation, epigenetic regulation and genome stability.^11, 12, 13, 14^ In addition, CTCF is a haploinsufficient tumour suppressor gene, which is essential for cell growth;^15, 16^ therefore, this disruption of CTCF function may contribute to tumour development and progression. In endometrioid carcinoma, *CTCF* is recurrently mutated, with more than 200 somatic mutations identified. The majority are inactivating mutations (71%), including 45% frameshift, 20% nonsense and 6% splice site mutations, while the remaining 29% are missense mutations enriched within the 11-ZF DNA binding domain, particularly in ZF4.^17, 18^ Importantly, these mutations can result in CTCF haploinsufficiency, but not complete loss, as CTCF is essential for somatic cell function.^17^ In contrast, *CTCF* is not commonly mutated in serous subtype carcinoma, but its function is primarily disrupted by copy number loss,^17^ indicating distinct mechanisms of CTCF functional impairment across histological subtypes. Genetic alterations of CTCF have been demonstrated to have a pro-tumorigenic effect in endometrial cancer via promoting cell survival and altering cell polarity.^17^

Given the essential role of CTCF in regulating gene expression, defining the molecular consequences of *CTCF* mutation in endometrial cancer is important. Our previous study identified extensive transcriptional dysregulation following sustained *CTCF* depletion in KLE EC cells.^17^ Among the dysregulated genes, *ZNF185*, which encodes an actin cytoskeleton-associated zinc finger protein, emerged as a candidate gene of interest, which was examined in this study. The aberrant expression of ZNF185 has been reported across various malignancies, including marked downregulation in prostate cancer, cervical, oesophageal, and head and neck squamous cell carcinoma,^19, 20, 21, 22^ but significant upregulation in pancreatic ductal carcinoma,^23^ suggesting that ZNF185 can exhibit both tumour-suppressing and tumour-promoting effects in different tumour types. However, its role in endometrial cancer and the mechanisms underlying its dysregulation in the context of *CTCF* loss have not been investigated.

In this study, we examined the effects of CTCF haploinsufficiency on ZNF185 expression in endometrial cancer cell lines using inducible shRNA knockdown. We identified a previously uncharacterised isoform of ZNF185 lacking an actin-targeting domain, designated ZNF185B and arising from an alternative promoter, and confirmed its upregulation following CTCF depletion in EC cells. In addition, we showed that the full-length and alternative isoform of ZNF185 exhibited distinct subcellular localisation, consistent with their ability to associate with actin. These findings were complemented with rapid degron-mediated depletion of CTCF in HEK293T cells showing robust induction of ZNF185B expression. Finally, we revealed that shRNA-mediated knockdown of ZNF185 expression reduced cell proliferation in endometrial cancer cells, suggesting an important role in tumour growth and progression.

## Results

### *CTCF* knockdown induces ZNF185 and APOE expression in KLE cells

As most *CTCF* mutations reported in endometrial carcinoma result in haploinsufficiency, the absence of detectable *CTCF* mutations in the KLE cell line made it a suitable model for studying the molecular consequences of reduced CTCF dosage. Our previous gene expression study using doxycycline (dox)-induced *CTCF* shRNA knockdown in KLE cells (**Supplementary Figure 1A & B**) revealed 1,744 differentially expressed genes by microarray analysis.^17^ The top downregulated and upregulated genes, with individual probe intensities highlighted for each locus are shown with *CTCF* one of the top downregulated genes as expected (**Figure 1A & B**). To validate these findings, selected differentially expressed genes were examined by qRT-PCR. *QDPR*, *TAMM41*, *C19ORF60* and *EPDR1* expression was downregulated upon dox-induced CTCF depletion compared to a non-targeting shRNA control, while *CAMK2B* and *ZNF185* were upregulated (all *p*<0.05, **Figure 1C & D**). We focused further on *ZNF185* as it encodes an actin-binding protein involved in cytoskeletal remodelling, and we previously observed a loss of cell polarity in KLE spheroids after *CTCF* knockdown.^17^ We also examined *APOE* encoding Apolipoprotein E due to its link to obesity, which is a known risk factor in endometrial cancer. At the protein level, western blot analysis validated the efficient knockdown of CTCF in KLE cells after 4 days of dox induction. The expression of ZNF185 (*p*=0.0315 and *p*=0.0062) and APOE (*p*=0.0496) were significantly increased upon CTCF depletion. Notably, two distinct ZNF185 protein species were detected using a ‘universal’ antibody which detects all ZNF185 isoforms (**Figure 1E**). The top bands are consistent with full-length ZNF185 protein (73.5 kDa) which is hereafter referred to as ‘ZNF185A’. The bottom band represents a smaller, uncharacterised ZNF185 isoform (∼35.0 kDa), which we designated as ‘ZNF185B’. The expression of both ZNF185 isoforms was significantly increased upon CTCF knockdown, with ZNF185B exhibiting a robust increase (> 2-fold, *p*=0.0062) (**Figure 1F**). We also performed the study in Ishikawa cells subjected to dox-induced *CTCF* shRNA knockdown and observed similar results (**Supplementary Figure 1B & C**). CTCF was robustly depleted after 4 days and led to concomitant induction of the ZNF185B protein isoform. Together, these data prompt further investigation of ZNF185 isoforms and their regulatory relationship with CTCF.

**Figure 1.**
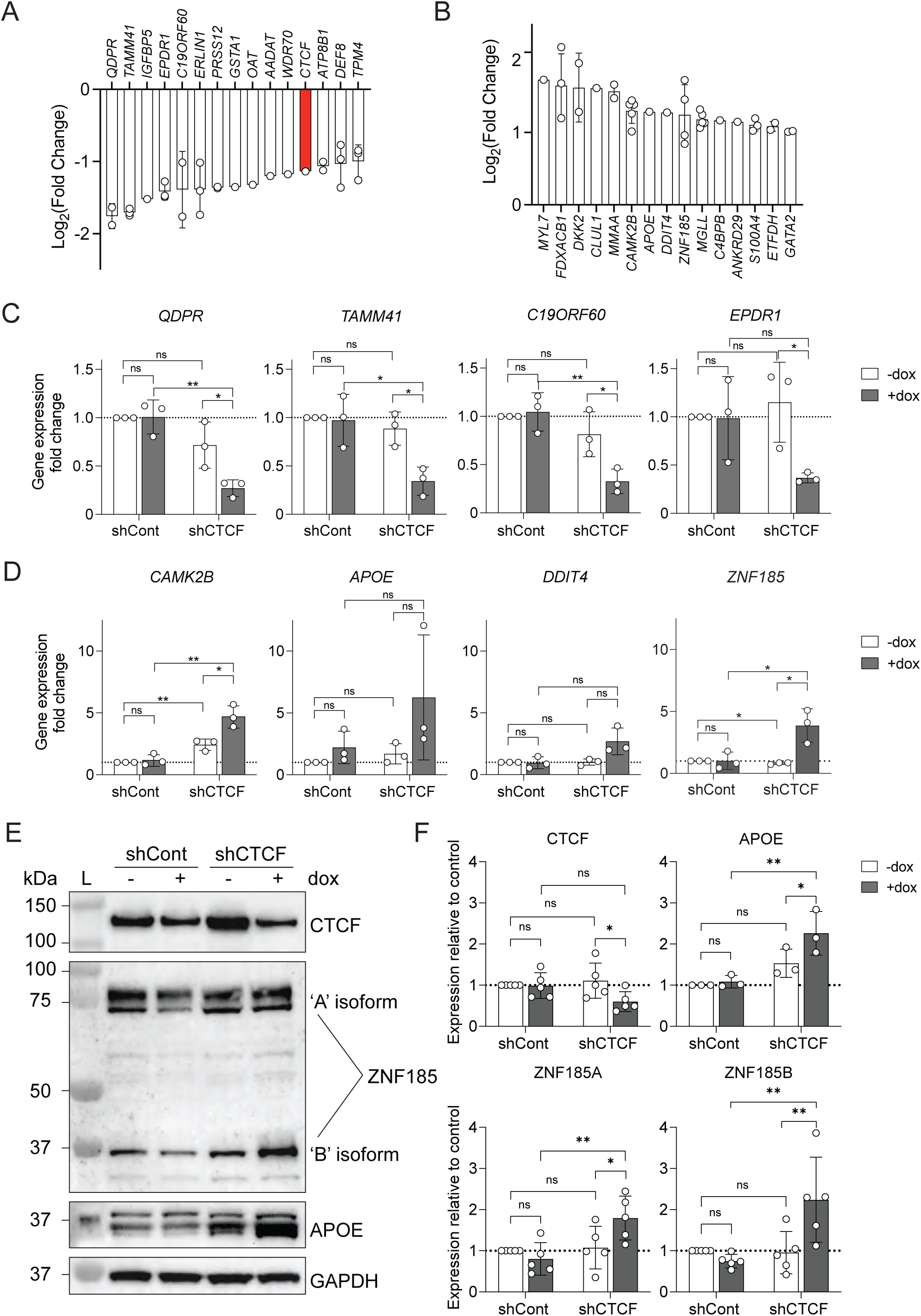
Activation of *ZNF185* and *APOE* expression after shRNA-mediated knockdown of *CTCF* in KLE cells. **A-B**) Summary data from a gene expression array following doxycycline-inducible *CTCF* knockdown for 4 days in KLE cells.^17^ A *CTCF*-targeting shRNA as well as a non-targeting control shRNA were used. The top down– **A**) and up-regulated genes **B**) with gene expression from individual probes at specific gene loci indicated as data points. Data shows the mean ± SD for n=1 to 5 datapoints. **C-D**) Validation of differentially expressed genes following doxycycline-inducible *CTCF* knockdown in KLE cells by qRT-PCR. Bar graphs for down-regulated genes **C**) and up-regulated genes **D**) are shown; data represents the mean ± SD of n=3 independent experiments each performed in triplicate. Statistical analysis was performed using Students t-test with *=*p*<0.05, **=*p*<0.01, ns=not significant. **E**) Representative western blot of KLE cell lysates after *CTCF* shRNA knockdown for 4 days which was probed with antibodies for CTCF, ZNF185 and APOE. GAPDH was used as a loading control, size markers (ladder, L) are indicated in kDa. **F**) Densitometric analysis of western blots from 3 independent experiments. Data represents the mean ± SD with statistical significance determined using 2-way ANOVA with Šídák’s multiple comparisons test with *=*p*<0.05, **=*p*<0.01, ns=not significant.

### Molecular characterization of ZNF185B isoform in endometrial cancer cell lines

To understand the molecular determinants of *ZNF185* isoforms, a schematic of the genomic locus of human *ZNF185* at chromosomal location Xq28 is shown (**Figure 2A**). The reference *ZNF185* transcript, or *ZNF185A*, originates from the transcription start site (TSS) within the canonical promoter (P1), whereas the novel isoform *ZNF185B* likely arises from a TSS within an alternative promoter (P2) ∼43 kb downstream. Importantly, both TSSs are located within CpG islands, and experimentally determined CTCF binding sites are dispersed throughout the locus. Notably, our previous microarray-based study could not distinguish between ZNF185 ‘A’ and ‘B’ isoform as the original probes used for previous gene expression microarray only detect 3’ regions within *ZNF185* mRNA (**Figure 2A**). These distinct transcripts may give rise to protein isoforms with different functional attributes. Both isoforms contain a C-terminal conserved *Lin-11*, *Isl-1* and *Mec-3* (LIM) domain, which facilitates protein-protein interactions in a range of biological processes including cytoskeletal organisation, cell signalling, cell migration and gene regulation (**Figure 2B**).^24, 25^ An N-terminal actin-targeting domain (ATD, 1-135 aa) is present in ZNF185A and is essential for mediating intracellular targeting, F-actin binding and cytoskeletal organisation.^19^ In contrast, ZNF185B lacks the ATD, but instead encodes a unique 41 aa N-terminus (**Figure 2B**). The lack of ATD may alter the association of ZNF185B with cytoskeletal actin structures and may affect cell polarity, subcellular interaction and cell proliferation.^19, 26, 27^

**Figure 2.**
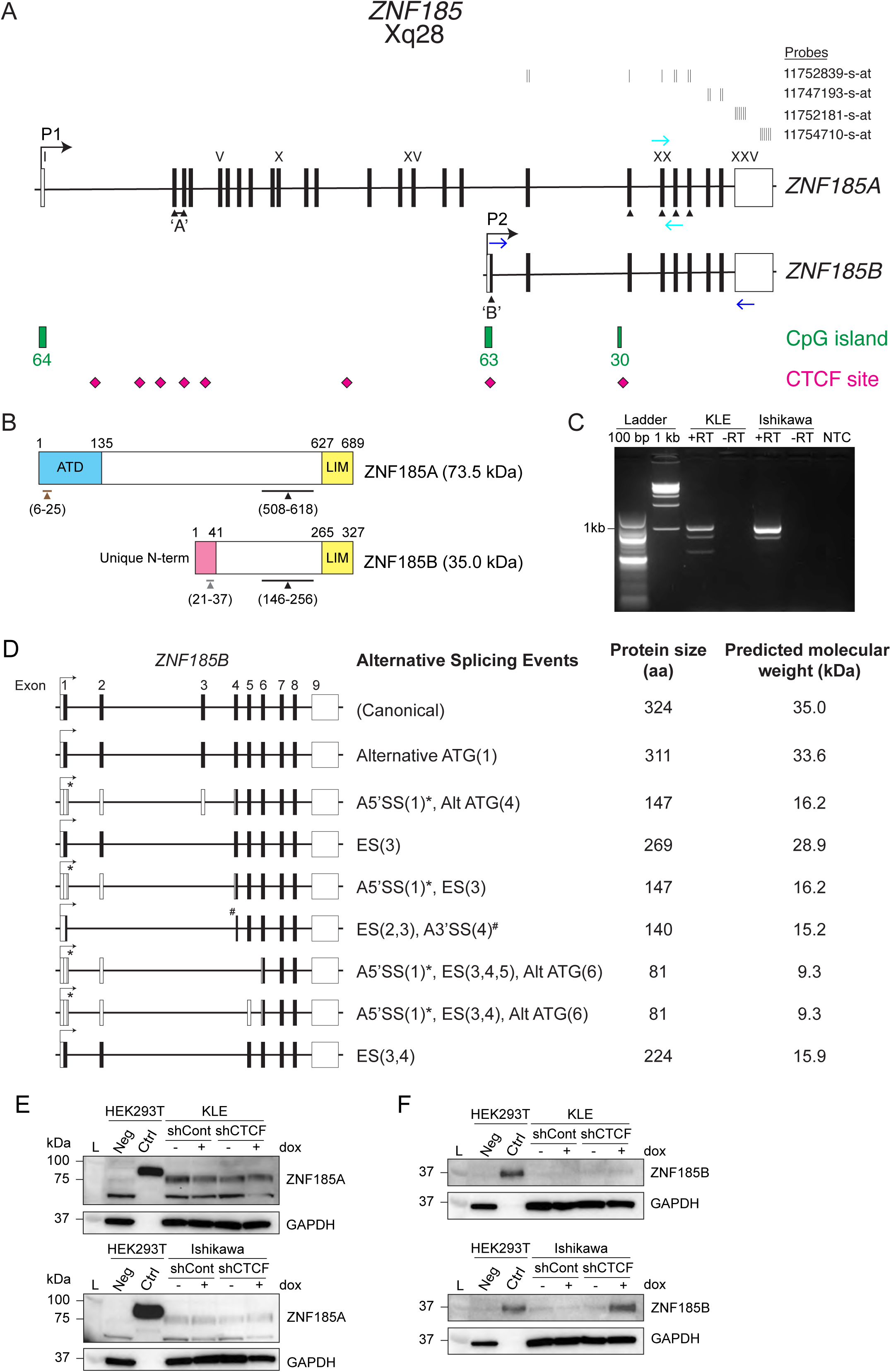
Molecular characterisation of *ZNF185* expression in KLE cells reveals a novel isoform, ZNF185B. **A**) Genomic locus of human *ZNF185* at chromosomal location Xq28. Coding (black boxes) and non-coding exons (empty boxes) are depicted and numbered. The transcription start site (TSS) for *ZNF185*, the canonical isoform named ‘*ZNF185A*’, and an alternative TSS at a downstream promoter giving rise to an alternative isoform named ‘*ZNF185B*’ are shown. The location of 3’ microarray probes related to Figure 1A & B are indicated for *ZNF185*. CpG islands (green boxes) and number of CpGs, along with putative CTCF binding sites (pink diamonds) are depicted. PCR primers used for qRT-PCR of *ZNF185* gene expression analysis in Figure 1D and RT-PCR in Figure 2C (cyan and blue arrows respectively). **B**) Schematic of ZNF185 protein isoforms. ZNF185A has an actin-targeting domain (ATD) at the N-terminus and a LIM domain at the C-terminus, whereas ZNF185B has a unique 41 amino acid N-terminus and a LIM domain at the C-terminus. The black lines indicate the immunogen for the commercial ZNF185 antibody, the brown line indicates the peptide immunogen for the custom ZNF185A antibody, and the grey line indicates the peptide immunogen for the custom ZNF185B antibody. **C**) RT-PCR of *ZNF185B* in KLE and Ishikawa cells using primers (blue) shown in (A); 100 bp and 1 kb ladders are shown. Samples were prepared with (+) or without (-) reverse transcription (RT); Non-template control (NTC) and –RT controls were included to ensure the absence of genomic DNA contamination and non-specific amplification. **D**) Schematic of *ZNF185B* transcripts cloned from Ishikawa cells. A summary table presents the identified transcripts along with predicted protein sizes. A5’SS = alternative 5’ splice site, indicated with *; A3’SS = alternative 3’ splice site, indicated with #; ES = exon skipping; brackets indicate exon number. **E-F**) Western blot validation of custom ZNF185 antibodies in endometrial cancer cell lines against ZNF185A **E**), and ZNF185B **F**). As controls, HEK293T cells were mock-transfected or transfected with either pcDNA3.1-HA-ZNF185A or pcDNA3.1-ZNF185B plasmids, and western blots were performed alongside KLE and Ishikawa cell lysates with or without *CTCF* shRNA knockdown for 4 days. Transfected HEK293T cell lysates were diluted 1:50 or 1:200 in cell lysis buffer prior to loading to prevent signal saturation due to overexpression. Blots were probed with custom antibodies for ZNF185A and ZNF185B to assess antibody specificity and isoform detection. GAPDH was used as a loading control, size markers (ladder, L) are indicated in kDa.

To experimentally verify these findings, we used RT-PCR analysis in KLE and Ishikawa cell lines and detected multiple *ZNF185B* transcripts (**Figure 2C**). One major transcript was identified corresponding to a 984 bp open reading frame of *ZNF185B* giving rise to an expected 327 aa protein. Several alternatively spliced transcripts from Ishikawa cells were confirmed by sequencing analysis which revealed alternative 5’ and 3’ splice site usage, and exon skipping (**Figure 2D**). The predicted molecular weights for these alternative ZNF185B isoforms are provided but were not experimentally verified.

To confirm the protein expression of ZNF185 isoforms, custom antibodies were designed to specifically detect both major isoforms. The ZNF185A antibody was raised against a peptide representing residues 6 – 25 within the ATD and the ZNF185B antibody was raised against a peptide representing residues 21 – 37 within the unique N-terminus (**Figure 2B**). HEK293T cells transfected with ZNF185A (HA-tagged) or ZNF185B expression plasmids were used as positive controls to validate custom antibodies specificity and to confirm the detection of proteins at the expected molecular weights. Western blot analysis was performed alongside lysates from KLE or Ishikawa cells expressing dox-inducible shRNAs in which *CTCF* knockdown had been confirmed, and verified the expression of ZNF185A (**Figure 2E**) and ZNF185B isoforms (**Figure 2F**). Collectively, these findings reveal a novel isoform ZNF185B expressed in endometrial cancer cell lines, which lacks the ATD. We have also validated specific antibodies which may be used to detect different functional characteristics of ZNF185A and ZNF185B.

### ZNF185 isoforms exhibit distinct subcellular expression and localisation

We next examined CTCF and ZNF185 expression in five endometrial cancer cells (**Figure 3A**) in which we previously confirmed *CTCF* mutation status.^17^ Notably, a nonsense G19* mutation in the first coding exon of *CTCF* in HEC1B, results in a truncated CTCF protein arising from an alternative downstream ATG. Using a universal ZNF185 antibody, ZNF185A was detected in all five cell lines, with higher expression in HEC1A and HEC1B and lower in KLE, Ishikawa and RL95-2 cells. A similar pattern was observed using ZNF185A-specific antibody, whereas the ZNF185B-specific antibody detected the ZNF185B isoform at a low level (**Figure 3A**).

**Figure 3.**
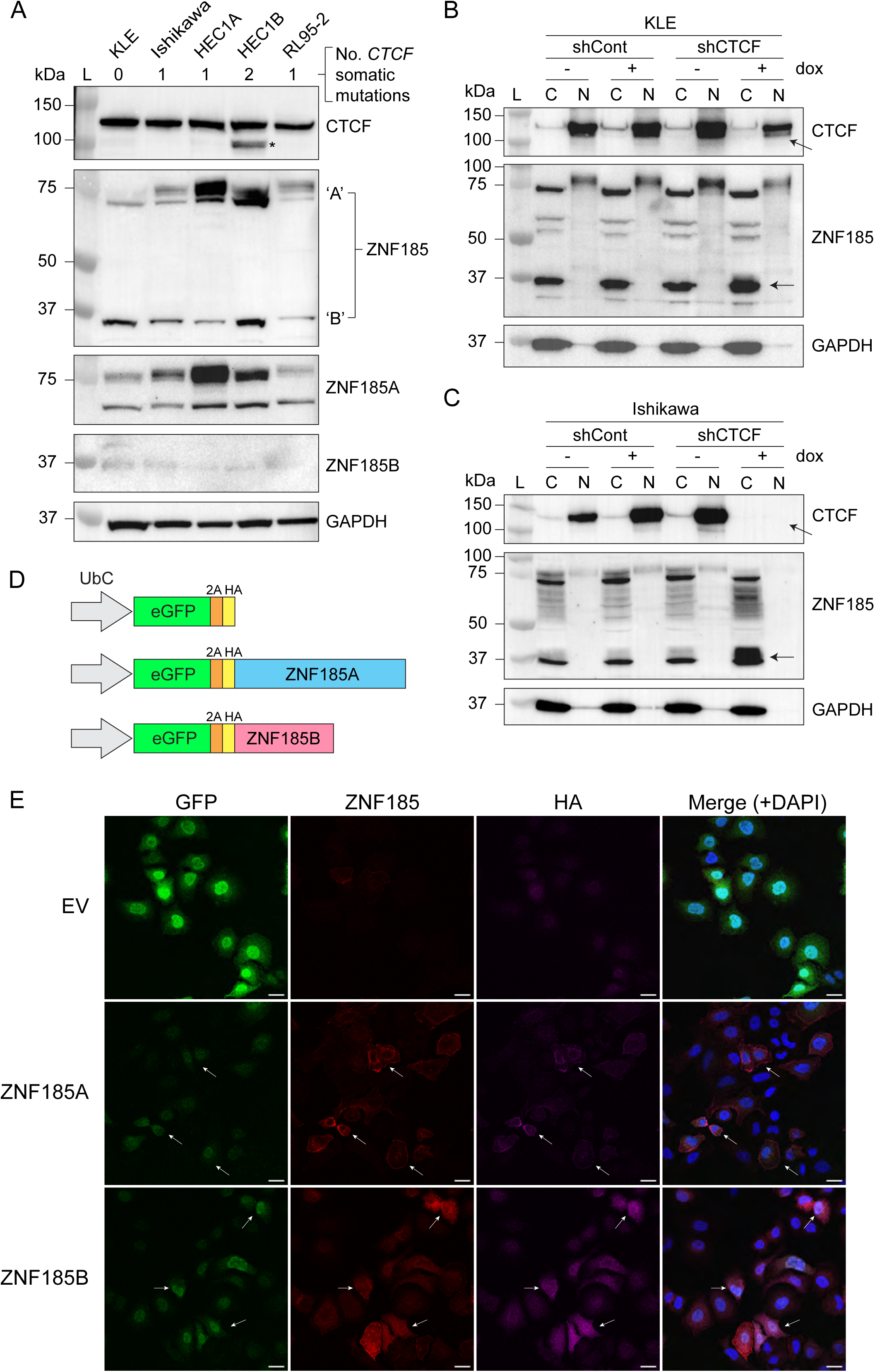
Distinct subcellular localisation observed for ZNF185 isoforms in endometrial cancer cells. **A**) Western blot of whole-cell lysates from five endometrial cancer cell lines probed with antibodies for CTCF, ZNF185, ZNF185A and ZNF185B isoforms. GAPDH was used as a loading control, and molecular weight markers (ladder, L) are indicated in kDa. The number of known somatic *CTCF* mutations in each cell line are also indicated; * indicates *CTCF* G19* mutation leading to alternative downstream ATG usage. **B-C**) Western blots of cytoplasmic (C) and nuclear (N) fractions from KLE **B**) and Ishikawa **C**) cells which were probed with antibodies against CTCF and ZNF185 to examine subcellular localisation. GAPDH was used as a loading control, the molecular weight ladder (L) is indicated. Arrowheads indicate reduced CTCF expression in the nucleus and increased ZNF185B expression in the cytoplasm in shCTCF cells after 4 days of dox treatment. **D**) Schematic of lentiviral constructs used to overexpress ZNF185A and ZNF185B isoforms. **E**) Immunofluorescence of Ishikawa cells transduced with lentivectors expressing empty vector (EV) or with ZNF185A or ZNF185B. Cells were stained with antibodies against ZNF185 (red) and HA (magenta). Transduced cells were GFP-positive (green), and nuclei were stained with DAPI (blue) in merged image. Arrows indicate representative transduced cells overexpressing ZNF185A or ZNF185B. Images were acquired with a 40× water immersion objective, scale bars = 25 µm.

To determine the subcellular distribution of ZNF185 in endometrial cancer cell lines, cytoplasmic and nuclear fractions were isolated from KLE and Ishikawa cells following 4 days of dox-induced shRNA knockdown of *CTCF*. Western blot analysis confirmed efficient extraction of nuclear and cytoplasmic proteins after probing for CTCF and GAPDH respectively (**Figure 3B & C**). ZNF185A was detected in both nuclear and cytoplasmic fractions with a higher molecular weight form present only in the nucleus in KLE cells and in both fractions in Ishikawa cells. In contrast, ZNF185B was only observed in the cytoplasmic fraction. Notably, after *CTCF* knockdown in both cell lines, a reduction in CTCF expression (arrows) was accompanied by a notable increase in cytoplasmic ZNF185B expression (arrows, **Figure 3B & C**), which suggests that ZNF185B is negatively regulated by CTCF.

To further validate the isoform-specific localisation of ZNF185, Ishikawa cells were transduced with lentiviral constructs containing HA-tagged ZNF185A, ZNF185B or an empty vector along with an eGFP reporter (**Figure 3D**). Successful transduction was examined by flow cytometry, and transduction efficiency was defined as the proportion of eGFP-positive cells (**Supplementary Figure 2A**). Western blot analysis of whole cell lysates confirmed successful overexpression at the protein level (**Supplementary Figure 2B**). The universal ZNF185 antibody and custom isoform-specific antibodies detected both endogenous ZNF185 and the overexpressed proteins, whereas the HA antibody only detected ectopic expression. We then examined the subcellular localisation of HA-tagged ZNF185A and ZNF185B in Ishikawa cells by immunofluorescence (IF) staining and confocal microscopy (**Figure 3E**). GFP fluorescence was used to identify transduced cells, and HA immunostaining demonstrated strong co-localisation with ZNF185 stained using the universal ZNF185 antibody. The results showed that ZNF185A was primarily enriched at the cell periphery, whereas ZNF185B exhibited a diffuse cytoplasmic distribution (arrows, **Figure 3E**). Similar patterns of staining were observed in HA-ZNF185A– and HA-ZNF185B-expressing cells stained with the custom ZNF185A and ZNF185B antibodies respectively (arrows, **Supplementary Figure 2C**). We further investigated whether ZNF185 associates with F-actin using phalloidin staining. Co-localisation between HA-tagged ZNF185A and F-actin was observed in certain cells (**Supplementary Figure 2C**), confirming actin association as expected. In contrast, HA-tagged ZNF185B did not exhibit co-localisation with F-actin, consistent with the lack of an ATD.

### Dose-dependent upregulation of ZNF185B following *CTCF* depletion

To examine the interrelationship between CTCF dosage and ZNF185B expression level in endometrial cancer cells, a time-course of doxycycline-inducible knockdown of CTCF was performed in KLE and Ishikawa cell lines over 4 days (**Figure 4A & C**). Efficient knockdown of CTCF was confirmed in both cell lines, with a modest (∼30%) reduction in KLE cells and a more robust (∼80%) reduction in Ishikawa cells (**Figure 4B & D**). With progressive CTCF depletion over time, ZNF185A expression remained largely unchanged in both cell lines, with only minor increases at selected time points, suggesting it was not responsive to CTCF depletion (**Figure 4B & D**). In contrast, ZNF185B exhibited a time-dependent increase following CTCF knockdown. Quantification of relative protein expression, defined as the ratio of shCTCF to shCont across time points and normalised to the 0 h time point, highlighted this phenomenon (**Figure 4B & D**). The increase became evident at 72 h in both cell lines and further increased over time, suggesting a cumulative effect. By 96 h, ZNF185B protein levels were approximately 3.0-fold higher in KLE cells and 2.8-fold higher in Ishikawa cells relative to the control conditions. Overall, these data suggest that CTCF negatively regulates ZNF185B expression in endometrial cancer cells in a dose-dependent manner. Importantly, this was a consistent observation irrespective of CTCF knockdown efficiency or cell line.

**Figure 4.**
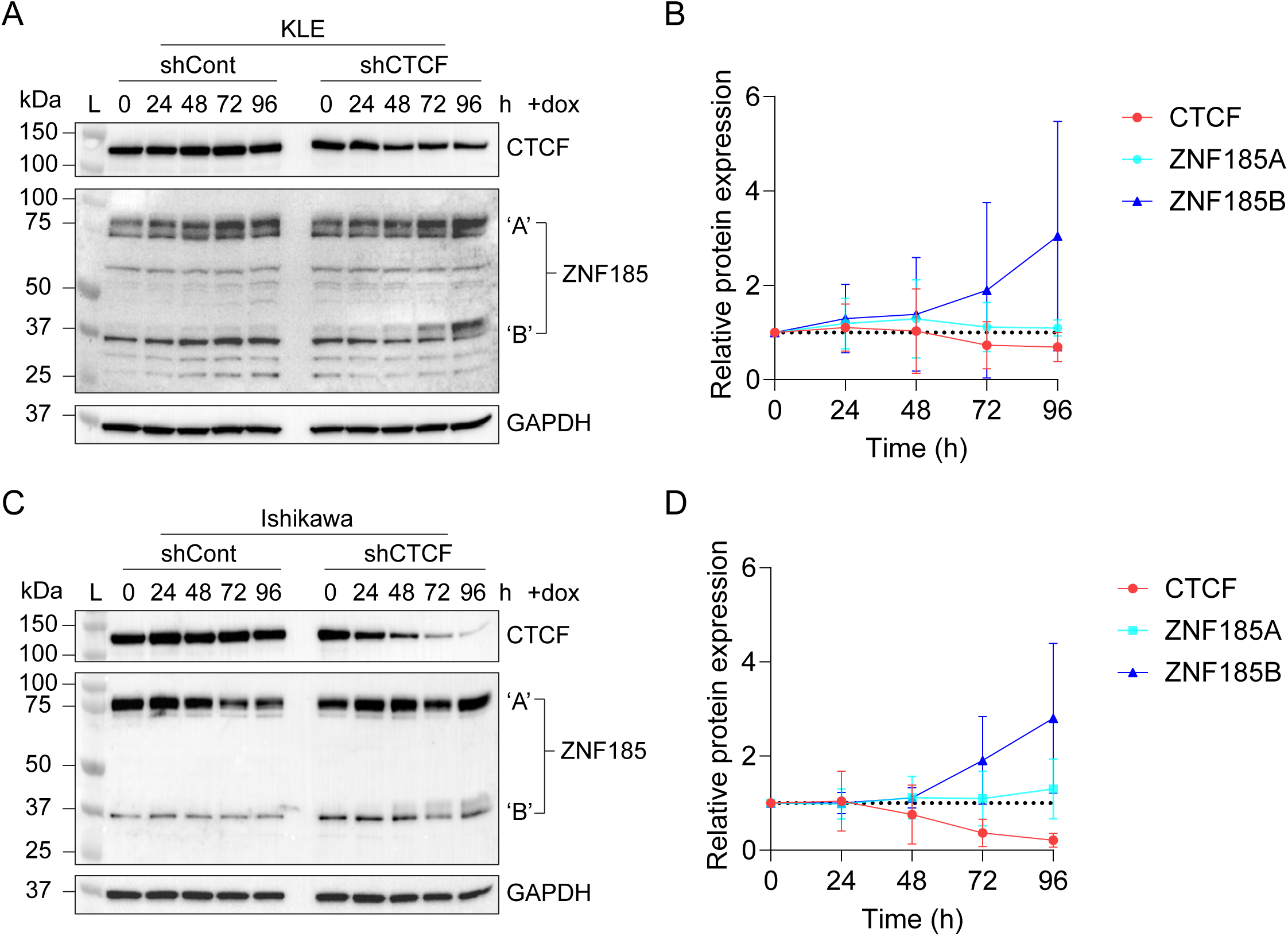
CTCF protein dosage regulates ZNF185B expression in endometrial cancer cells. Representative western blots of KLE **A**) or Ishikawa **C**) cell lysates collected at 0, 24, 48, 72, and 96 h following *CTCF* shRNA knockdown (1 µg/mL dox) which was probed with antibodies for CTCF and ZNF185. GAPDH was used as a loading control, size markers (ladder, L) are indicated in kDa. **B, D**) Densitometric analysis of western blots, data represents the mean ± SD from 3 independent experiments. Protein expression was first normalised to GAPDH, then the relative protein expression was defined as the ratio of protein levels in shCTCF cells to the corresponding shCont cells at each time point, with values further normalised to 0 h.

### Acute auxin-mediated depletion of *CTCF* induces ZNF185B expression

To study the effect of rapid CTCF depletion, an auxin-inducible degron (AID) for CTCF was established in HEK293T cells.^28^ Firstly, this required the stable integration of an auxin receptor AtAFB2 into the adeno-associated virus integration site 1 (*AAVS1*) locus in HEK293T cells via CRISPR-Cas9-mediated homology-directed repair (HDR). Secondly, the HEK293T-AtAFB2 cells were engineered to incorporate an AA7 auxin-inducible degron and 3x FLAG tag at the C-terminus of the endogenous *CTCF* gene via CRISPR-Cas9-mediated HDR, thereby enabling auxin-inducible depletion of *CTCF* (**Figure 5A**).^28^ Two independent clones, 4.22 and 4.24, with confirmed knock-in of AA7-3FLAG on all alleles were obtained. Western blot analysis was performed and demonstrated effective acute depletion of CTCF in both clones, with a robust reduction observed after 1 h of indole-3-acetic acid (IAA) treatment (**Figure 5B**), which were also confirmed by densitometric analysis (**Figure 5C**). Following this, immunoblotting using custom ZNF185A and ZNF185B antibodies revealed a small increase in ZNF185A expression at a later timepoint, whereas a robust increase in ZNF185B expression was observed from 24 h of IAA treatment, reaching 4.6-fold in clone 4.22 and 11.5-fold in clone 4.24 at 48 h compared to untreated cells.

**Figure 5.**
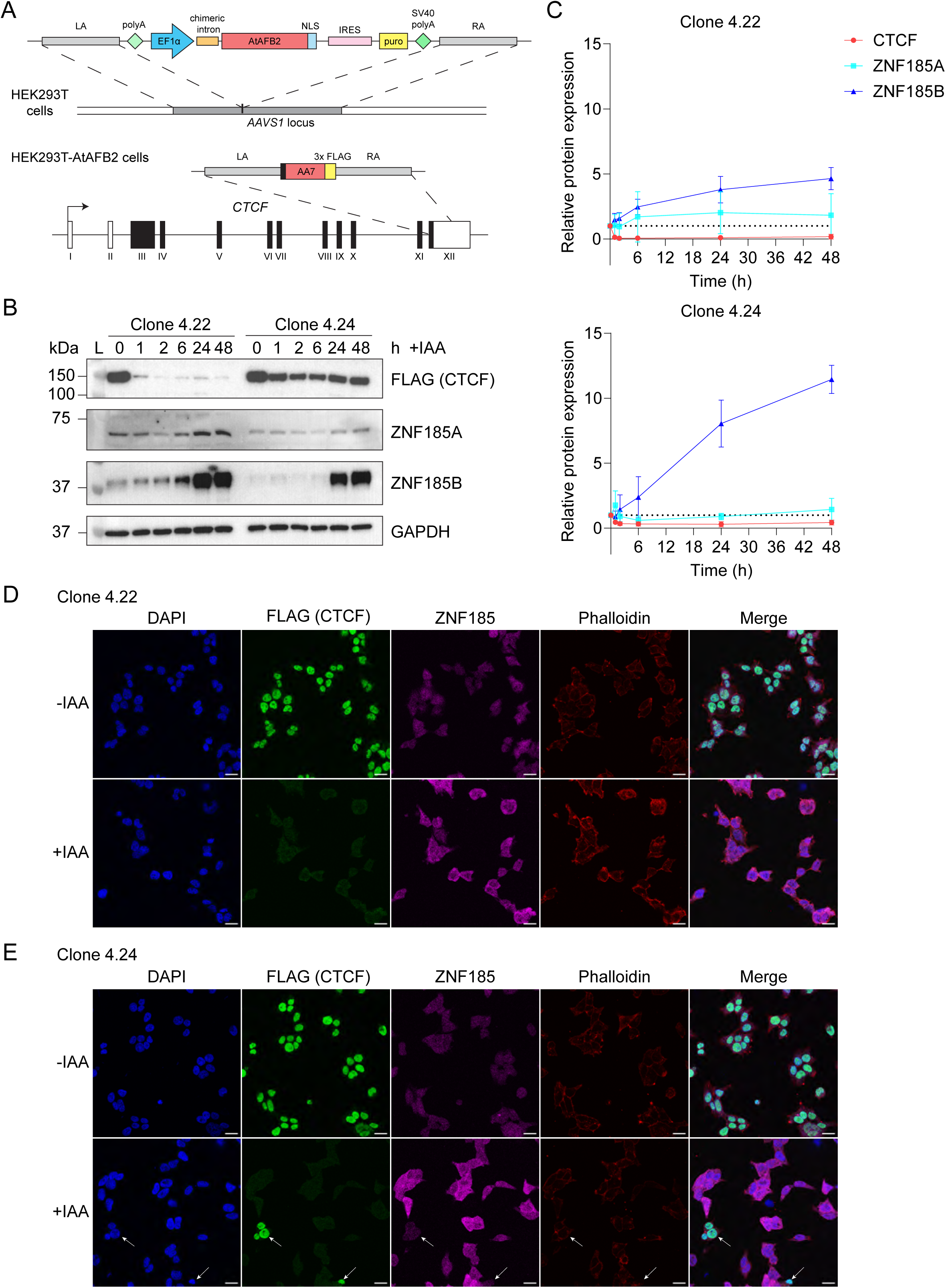
Upregulation of ZNF185B upon rapid auxin-mediated degradation of CTCF. **A**) Schematic of the *CTCF* locus targeted with AID in HEK293T cells. **B**) Western blot of whole cell lysates from two CTCF-AID clones established in HEK293T cells collected at 0, 1, 6, 12, 24 and 48 h following indole-3-acetic acid (IAA) treatment were probed with antibodies for FLAG (CTCF) as well as ZNF185A and ZNF185B isoforms. GAPDH was used as a loading control, the molecular weight ladder (L) is indicated. **C**) Densitometric analysis of Western blots, data represents the mean ± SD from 3 independent experiments. Protein expression levels were normalised to GAPDH and the 0 h time point. **D–E**) Immunofluorescence of Clone 4.22 **D**) and Clone 4.24 **E**) cells following IAA treatment. Cells were stained for FLAG (CTCF, green), ZNF185 (magenta), and phalloidin (for F-actin, red), with nuclei stained with DAPI (blue). Images were acquired using a 40× water immersion objective, scale bars = 25 µm. Arrows indicate cells that didn’t undergo auxin-mediated CTCF depletion.

We further confirmed the upregulation of ZNF185 following *CTCF* depletion by IF, which showed enhanced intensity of diffuse ZNF185 staining following IAA treatment in both clones (**Figure 5D & E**). Notably, CTCF was nearly undetectable in clone 4.22 (**Figure 5D**), whereas in clone 4.24, some cells appeared resistant to *CTCF* depletion and did not exhibit corresponding increased expression of ZNF185 (arrows, **Figure 5E**). This immunostaining pattern was consistent with the different knockdown efficiency observed in western blot analysis. Altogether, these findings indicate that acute depletion of CTCF induces ZNF185 expression, particularly the ZNF185B isoform, suggesting a broader role of CTCF in regulating ZNF185 expression beyond endometrial cancer cell lines.

### Depletion of ZNF185 expression inhibits endometrial cancer cell proliferation

To examine the effects of ZNF185 on endometrial cancer cell growth, three independent shRNAs were used to knockdown *ZNF185* expression in KLE and Ishikawa cells (**Supplementary Figure 3A)**. Western blot analysis and densitometric quantification confirmed a predominant knockdown of ZNF185A in KLE cells, whereas ZNF185B was not affected (**Figure 6A & C**). However, in Ishikawa cells, both ZNF185A and ZNF185B were efficiently depleted (**Figure 6B & D**). After stable knockdown of ZNF185 in both endometrial cancer cell lines, MTT assays were performed to assess cell growth. In KLE cells, ZNF185 depletion resulted in a significant reduction in cell proliferation over 8 days, as indicated by consistently lower absorbance across all three independent shRNAs compared to the non-targeting control (shCont) (**Figure 6E**). A similar inhibitory effect on cell growth was also observed in Ishikawa cells following ZNF185 knockdown over 4 days (**Figure 6F**). In addition, clonogenicity assays were performed to assess colony forming ability. Colony number was significantly reduced in both KLE and Ishikawa cells following ZNF185 knockdown (**Figure 6G & H & Supplementary Figure 3B**), indicating an impaired long-term proliferative capacity of cancer cells upon ZNF185 depletion. In contrast, the overexpression of either ZNF185A or ZNF185B had minimal impact on cell growth or colony formation in KLE cells (**Supplementary Figure 3C & D & E**). Taken together, these findings demonstrated that ZNF185 knockdown leads to impaired cell growth in endometrial cancer cells.

**Figure 6.**
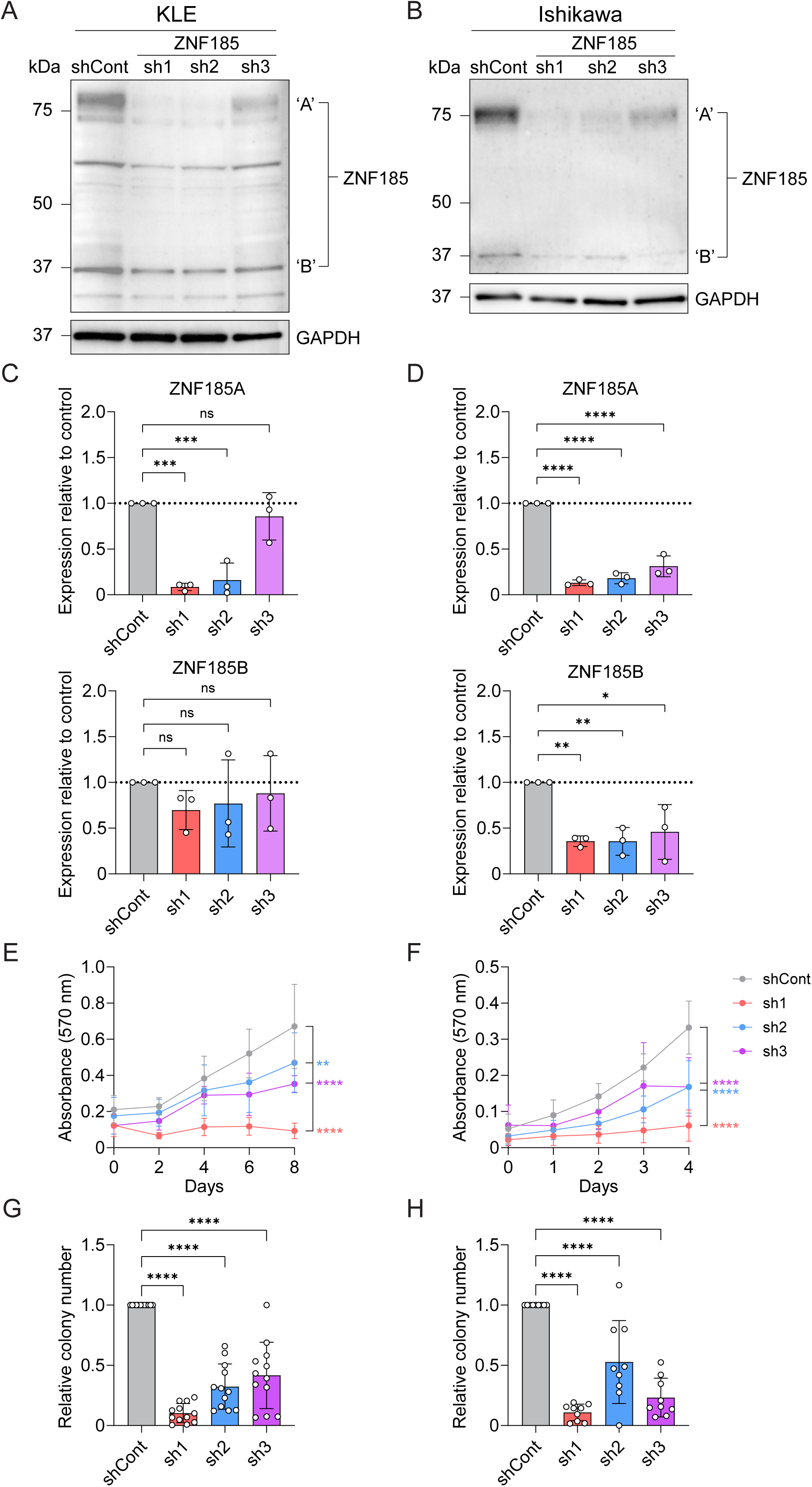
Effects of ZNF185 knockdown on endometrial cancer cell proliferation. **A-B**) Representative western blot of KLE cells **A**) or Ishikawa cells **B**) transduced with 3 independent shRNAs targeting human *ZNF185* mRNAs or a non-targeting control shRNA (shCont) were probed with antibodies against universal ZNF185 and CTCF. GAPDH was used as a loading control and molecular weight markers are indicated in kDa. **C-D**) Densitometric analysis of western blots of KLE **C**); and Ishikawa cells **D**) from 3 independent experiments. Data represents the mean ± SD with statistical significance determined using one-way ANOVA with Dunnett’s multiple comparisons test with *=*p*<0.05, **=*p*<0.01, ***=*p*<0.001, ****=*p*<0.0001, ns=not significant. **E-F**) MTT absorbance reading measured in KLE cells **E**) transduced with *ZNF185* shRNAs over 8 days of n=4 independent experiments each performed in triplicate; and in Ishikawa cells **F**) transduced with *ZNF185* shRNAs over 4 days of n=3 independent experiments each performed in triplicate. Data represents the mean ± SD with statistical significance determined at the last time point using one-way ANOVA with Dunnett’s multiple comparisons test with *=*p*<0.05, **=*p*<0.01, ***=*p*<0.001, ****=*p*<0.0001, ns=not significant. **G-H**) Clonogenicity assay of KLE cells **G**) transduced with ZNF185 shRNAs after 21 days incubation from 4 independent experiments each performed in triplicate; and Ishikawa cells **H**) transduced with ZNF185 shRNAs after 10 days incubation from 3 independent experiments each performed in triplicate. Colony number was normalised to shCont. Data represents the mean ± SD with statistical significance determined using one-way ANOVA with Dunnett’s multiple comparisons test with *=*p*<0.05, **=*p*<0.01, ***=*p*<0.001, ****=*p*<0.0001, ns=not significant.

## Discussion

Clinicopathological studies have revealed an association of ZNF185 expression with liver metastasis in colon cancer and hematogenous metastasis in pancreatic ductal adenocarcinoma (PDAC) patients.^23, 29^ Furthermore, increased venous invasion was observed with plasma membrane-specific ZNF185 staining in PDAC. However, in metastatic spread of oesophageal squamous cell carcinoma, ZNF185 is significantly suppressed in lymph node metastasis.^30^ These seemingly disparate findings illustrate that the molecular contribution of ZNF185 to cancer growth and spread is poorly understood.

Our previous study revealed that knockdown of CTCF expression in KLE endometrial cancer cell-derived organoids led to cellular disorganisation due to loss of cell polarity.^17^ ZNF185 as a significantly upregulated gene was investigated due to its known association with actin and involvement in cytoskeletal organisation. We confirmed this ZNF185 upregulation upon depletion of CTCF as well as the presence of a previously uncharacterised isoform, ZNF185B, lacking the key ATD using available and custom antibody reagents. We further demonstrated that ZNF185B expression was negatively regulated by CTCF in a dose-dependent manner, a regulatory relationship observed in two endometrial cancer cell lines. A similar observation was observed in an acute CTCF depletion degron model supporting a direct role for CTCF protein expression levels in tuning ZNF185 expression.

The co-location of the canonical *ZNF185* promoter (P1)^21^ and an alternative promoter (P2)^21^ ∼43.0 kb downstream within CpG islands and proximal to *CTCF* binding sites, suggests that *ZNF185* expression may be regulated by both epigenetic and *CTCF*-driven mechanisms. An alternative transcription start site juxtaposed in the middle of the *ZNF185* locus and giving rise to protein isoforms of different lengths and lacking key functional domains, is reminiscent of transcription factor *ELF2*, in which we identified a similar genomic structure and molecular ontogeny of ‘A’ and ‘B’ isoforms.^31^ Custom isoform-specific antibodies were able to verify the presence of these ZNF185 isoforms, but also provide key evidence for different subcellular localisation and expression levels particularly upon depletion of CTCF.

The distinct subcellular localisation of ZNF185 isoforms is likely due to their structural differences. ZNF185A contains an N-terminal ATD and C-terminal LIM domain, whereas ZNF185B lacks the ATD at the N-terminus but retains the C-terminal LIM domain. As LIM domain containing proteins can localise to both the nucleus and cytoplasm,^25^ this was consistent with our western blot findings showing ZNF185A presence in both compartments. In addition, ZNF185 directly binds to F-actin via its ATD, rather than LIM domain.^19^ This explains our observation of cell cortical localisation of ZNF185A and cytoplasmic distribution of ZNF185B. However, the absence of ATD does not exclude a possible role of ZNF185B in cytoskeletal regulation, as it may instead participate indirectly through LIM domain-mediated interactions or other signalling pathways, such as protein kinase A (PKA)/RhoA signalling, whereby ZNF185 is associated with inhibition of RhoA activity.^27^

As loss of CTCF disrupts cell polarity in KLE spheroids,^17^ the upregulation of ZNF185 expression may also contribute to this process and directly influence epithelial-mesenchymal transition (EMT), a key feature of tumour progression. Although direct evidence linking ZNF185 to EMT is limited, its association with epithelial proteins suggests a potential involvement in regulating epithelial organisation. Notably, E-cadherin, an important component of adherens junctions and a classic epithelial EMT marker, is associated with endometrial cancer development and considered as a putative marker for good prognosis.^32, 33^ E-cadherin was shown to interact with, as well as co-localise with ZNF185, with their expression levels positively correlated in squamous cell carcinoma.^21^ Therefore, dysregulation of ZNF185 may perturb EMT-related processes, thereby contributing to the disruption of cell polarity.

Increasing evidence shows that ZNF185 plays an important role in regulating cell proliferation. In prostate cancer, overexpression of ZNF185 suppressed cell proliferation and anchorage-independent growth.^19^ Similarly, ZNF185 overexpression also reduced cell proliferation and invasion in lung adenocarcinoma cells.^34^ Consistent with these findings, ZNF185 knockdown enhanced the proliferation of oesophageal cancer cells.^22^ However, in this study, we observed that ZNF185 depletion reduced both endometrial cancer cell proliferation and clonogenic potential. This indicates that ZNF185 expression maybe required for endometrial cancer cell proliferation which has implications for endometrial tumour pathophysiology. In contrast, overexpression of ZNF185 had little effect on endometrial cancer cell growth and proliferation, suggesting enforced overexpression does not further induce cell proliferation.

In conclusion, we have identified a novel isoform of ZNF185, ZNF185B, that is dysregulated in *CTCF*-depleted endometrial cancer cells. Although ZNF185 has been reported to function as a tumour suppressor in several cancer types, our findings show that ZNF185 is upregulated by CTCF in a dose-dependent manner, and its loss inhibits cell proliferation, together suggesting a tumour-promoting role of ZNF185 in endometrial cancer cells in the context of *CTCF* haploinsufficiency.

## Material and Methods

### Cell culture

KLE, Ishikawa, HEC1A, HEC1B and RL95-2 endometrial cancer cell lines were generously gifted by Professor Deborah Marsh (University of Technology Sydney, Sydney, Australia) and HEK293T cells were obtained from St. Jude Children’s Research Hospital. All endometrial cancer cell lines were cultured in DMEM/F12 medium (Gibco, USA) supplemented with 10% (v/v) fetal bovine serum (FBS) and 1% (v/v) penicillin-streptomycin at 37 °C and 5% CO_2_ with or without 1 μg/mL doxycycline (dox, Sigma-Aldrich, USA) as indicated. HEK293T cells were cultured in DMEM medium (Gibco, USA) supplemented with 10% (v/v) FBS and 1% (v/v) penicillin-streptomycin at 37 °C and 5% CO_2_ with or without 100 μg/mL indole-3-acetic acid (IAA, Sigma-Aldrich, USA) as required.

### Vectors for gene expression studies

Dox-inducible pFH1tUTG-based vectors, containing a tetracycline (Tet)-responsive H1 promoter for shRNA expression, a ubiquitin-C promoter for the expression of a tetR-T2A-eGFP cassette, were engineered to express shRNAs targeting human *CTCF* or an *Arabidopsis thaliana* mir159a.^17^ For *ZNF185* shRNA knockdown, the lentiviral vectors used were based on the pLKO.1 plasmid backbone containing a U6 promoter for shRNA expression and an eGFP-2A-puro cassette driven by a human phosphoglycerate kinase (pgk) promoter for marking and selecting transduced cells. Oligonucleotides for 3 independent shRNAs targeting human *ZNF185* mRNAs as well as a non-targeting control shRNA with complementary overlapping ends for *AgeI/EcoRI* restriction endonuclease sub-cloning into the lentivector were obtained (Integrated DNA Technologies). For ZNF185 isoform overexpression, a ZNF185 human cDNA (clone 100068255, Horizon Discovery) was used as a template for PCR to amplify ZNF185A for cloning into pcDNA3.1-HA for N-terminal tagging with a HA epitope. To generate a ZNF185B expression vector, a 570 bp geneblock (IDT) encoding residues 1-178 was cloned into pcDNA3.1-HA-ZNF185A with *EcoRI*/*XhoI* sites. Both HA-tagged ZNF185 cDNAs were then subcloned into pFUW-eGFP-2A lentiviral backbone with *EcoRI*/*XbaI* restriction endonucleases for simultaneous eGFP expression with ZNF185 isoforms. Sequences for all shRNA oligos and primers are provided in **Supplementary Table 1**.

### Lentiviral vector production

Lentiviral supernatant was generated by the calcium phosphate co-transfection of HEK293T cells with a lentivector together with packaging plasmids pRSV-Rev, pMD2-g/pRRE and pMD2-VSV-G. After 24 h, the medium was replaced with fresh DMEM supplemented with 2 mM sodium butyrate (Sigma-Aldrich, USA), and cells were incubated for a further 24 h. Viral supernatants were collected by filtration through a 0.45 μm filter and snap-frozen in liquid nitrogen. Cells (3-5 × 10^5^) were seeded in a 6-well plate and incubated overnight before transduction. Viral supernatant was added in appropriate medium supplemented with 4 μg/mL polybrene (Sigma-Aldrich, USA) and transferred onto cells. Cells were spinoculated for 1.5 h at 1,500 rpm and then incubated with lentivirus for 48-96 h. Cells transduced with non-targeting shRNA or empty-vector lentivirus were used as controls. Transduction efficiency was evaluated by flow cytometry (LSR-Fortessa, BD) and fluorescence-activated cell sorting (FACS) was applied when enrichment of transduced cells was required.

### Flow cytometry

Cells were collected 48-96 h after transduction, washed with cold FACS wash buffer (2% (v/v) FBS in PBS), and resuspended in FACS buffer containing 2 μg/mL DAPI. Samples were kept on ice until analysis using a BD LSR-Fortessa Cell Analyser (BD Biosciences, USA). Dead cells were excluded based on DAPI staining. Flow cytometry data were analysed using FlowJo software. For enrichment of transduced cells, FACS was performed to isolate GFP– and mCherry-positive cells as required. Cells were resuspended in cold sterile FACS buffer, filtered through FACS tubes and stained with DAPI (1:1,000). Samples were kept on ice prior to cell sorting which was performed by Sydney Cytometry Facility. Sorted cells were subsequently pelleted and resuspended in the corresponding culture medium for further growth.

### RT-PCR, qRT-PCR, and DNA sequencing

Total RNA was extracted from cells using Trizol (Invitrogen, USA) following the manufacturer’s protocol. Then, cDNA was synthesised from 1 μg total RNA using iScript gDNA Clear cDNA Synthesis Kit (Bio-Rad, USA). Gene targets were amplified from cDNA by PCR using specific primers (see **Supplementary Table 1**) under the following cycling conditions: initial denaturation at 98 °C for 30 s, followed by 35 cycles of 98 °C for 10 s, 60 °C for 20 s and 72 °C for 30 s, with a final extension at 72 °C for 10 min. PCR products were electrophoresed on a 1-2% (w/v) agarose gel containing GelRed Nucleic Acid Stain (Biotium, USA) and visualised under UV light. Target bands were excised, purified using a Gel and PCR Clean-up Kit (Macherey-Nagel, Germany) and cloned into pCR-Blunt II-TOPO Vector (Invitrogen, USA) according to the manufacturer’s protocol. Recombinant plasmids were transformed into competent *E. coli*, and positive colonies were selected by kanamycin selection. Plasmid DNA was isolated from selected clones using NucleoSpin Plasmid kit (Macherey-Nagel, Germany) and sent for DNA sequencing. Sequencing results were aligned to the reference transcript sequence to identify transcript structure and evaluate potential alternative splicing events.

For quantitative RT-PCR, cDNA was synthesised as described above. Reactions were prepared using SYBR Green master mix in 10 μL final volume containing diluted cDNA and 10 μM gene-specific primers (see **Supplementary Table 1**). Amplification was performed on a CFX96TM Real-Time System C1000 Thermal Cycler (Bio-Rad, USA) using the following program: 95 °C for 3 min, followed by 40 cycles of 95 °C for 15 s, 60 °C for 20 s and 72 °C for 20 s, with a final extension at 72 °C for 20 s. A melt curve analysis was performed at the end of the amplification to confirm the specificity of PCR products. Relative transcript levels were calculated using the 2^−ΔΔCt^ method and normalised to housekeeping gene *B2M*. ‘No template’ and ‘no reverse transcription’ controls were included to exclude contamination and genomic DNA amplification.

### Plasmid transfection

*ZNF185B* PCR products corresponding to the predicted size were excised from agarose gels, purified, and submitted for DNA sequencing. After sequence verification, nested PCR was performed to get the expected insert and then cloned into a pcDNA3.1-based plasmid. The resulting construct was confirmed by restriction enzyme digestion and sequencing. HEK293T cells were seeded into 6-well plate and transfected at 60-70% confluence with pcDNA3.1-C-HA-target plasmid using the calcium phosphate transfection method. Cells were collected and lysed at 48 h after transfection.

### Protein extraction and immunoblotting

Cells were washed with phosphate-buffered saline (PBS) and lysed in cell lysis buffer (50 mM Tris-HCl, pH 7.5-8; 150 mM NaCl; 1% (v/v) Triton X-100) supplemented with protease inhibitor cocktails to obtain total protein. Nuclear and cytoplasmic protein fractions were extracted from cells using NE-PER Nuclear and Cytoplasmic Extraction Kit (Thermo Fisher Scientific, USA) following the manufacturer’s instructions. Equal amounts of protein lysate were mixed with 4X loading buffer containing NuPAGE LDS Sample Buffer and 100 mM Dithiothreitol (DTT) and denatured at 80 °C for 10 min. Samples were electrophoresed by SDS-PAGE using BOLT 4-12% Bis-Tris Protein Gels (Thermo Fisher Scientific, USA) and transferred onto a PVDF membrane (Merck Millipore, Germany) in a semi-dry transfer cell (Bio-rad, USA). Membranes were blocked in 5% (w/v) skim milk in PBS with 0.1% (v/v) Tween-20 (PBST) for 30 min at room temperature. Membranes then were incubated overnight at 4 °C with primary antibodies: rabbit anti-CTCF (1:5,000, Cell Signalling Technology, USA, 3418S), rabbit anti-ZNF185 (1:2,500, Sigma-Aldrich, USA, HPA000400), rabbit anti-ZNF185A (1:5,000, custom rabbit polyclonal antibody raised against a peptide representing residues 6-25 of ZNF185 conjugated to keyhole limpet hemocyanin (Mimotopes, Australia)), rabbit anti-ZNF185B (1:5,000, custom rabbit polyclonal antibody raised against a peptide representing residues 21-37 of ZNF185B conjugated to keyhole limpet hemocyanin (Mimotopes)), goat anti-APOE (1:10,000, Sigma-Aldrich, USA, AB947) or mouse anti-GAPDH (1:5,000, Abcam, UK, AB8245). Membranes were washed 3 times for 5 min each in PBST and then incubated with donkey anti-mouse/rabbit HRP-conjugated (1:5,000, Merck Millipore, USA, AP192P, AP182P) or mouse anti-goat HRP-conjugated (1:5,000, Santa Cruz, USA, sc-2020) secondary antibodies for 1 h at room temperature. Alternatively, membranes were incubated with mouse anti-FLAG-HRP (1:5,000, Sigma-Aldrich, USA, A8592) for 2 h or mouse anti-HA-HRP (1:5,000, Cell Signalling Technology, USA, 2999S) for 3 h at room temperature. After three washes in PBST, blots were visualised using an enhanced chemiluminescent reagent and imaged with a Bio-Rad ChemiDoc imaging system. Densitometry analysis was performed using Image J software. Target band intensities were normalised to loading control, and values were expressed relative to the control.

### Immunofluorescence and imaging

Cells (4-8 × 10^4^) were seeded in a μ-Slide 8-well chamber slide (Ibidi, Germany) and incubated at 37 °C and 5% CO_2_ for overnight to allow adherence. Cells were rinsed in PBS on the following day and fixed in fresh 4% (w/v) formaldehyde (Thermo Fisher Scientific, USA) in PBS at room temperature for 15 min. Samples were washed 3 times for 5 min each in PBS and then permeabilised with freshly prepared 0.1% (v/v) Triton X-100 (Thermo Fisher Scientific, USA) in PBS for 10 min at room temperature. Samples were washed again in PBS as above, followed by blocking with 5% (v/v) goat serum (Gibco, USA) in 0.02% (v/v) Triton X-100 in PBS for 30 min at room temperature. Primary antibodies were diluted appropriately in the blocking buffer and applied to stain samples for 1.5 h at room temperature in the dark. Primary antibodies used included mouse anti-FLAG (1:100, Sigma-Aldrich, USA, F1804), mouse anti-HA (1:500, BioLegend, USA, 901501), rabbit anti-ZNF185 (1:100, Sigma-Aldrich, USA, HPA0004000), rabbit anti-ZNF185A (1:100, custom), rabbit anti-ZNF185B (1:100, custom). Samples were washed 3 times for 5 min each in PBS and then incubated with fluorescence-conjugated secondary antibodies including goat anti-mouse IgG Alexa Fluor 488 (1:500, Invitrogen, USA, A32723), goat anti-mouse IgG Alexa Fluor 594 (1:500, Invitrogen, USA, A11020), goat anti-rabbit IgG Alexa Fluor 594 (1:500, Invitrogen, USA, A11012), goat anti-rabbit IgG Alexa Fluor 647 (1:500, Molecular Probes, USA, A21245) and phalloidin-Alexa Fluor 647 (1:750, Invitrogen, USA, A30107) diluted in blocking buffer for 1 h at room temperature in the dark. Samples were washed with PBS containing 0.2 μg/mL DAPI to counterstain nuclei for 3 min and then washed twice with PBS. Samples were mounted with mounting media (Ibidi, Germany) and protected from light until imaging. Images were acquired with a Leica TCS SP8 confocal microscope using a 40X objective and processed using ImageJ.

### Establishment of auxin-inducible degron (AID) of CTCF

An auxin-inducible degron of *CTCF* was established with CRISPR-Cas9 homology-directed repair using a two-step method.^28^ To achieve this, pSH-EFIRES-P-AtAFB2-mCherry-weak-NLS (#129717, Addgene) was modified with a geneblock (Integrated DNA Technologies) cloned in with *BsrGI/NotI* to remove mCherry but keep the weak C-terminal NLS (GGAKRVKLD). Then, HEK293T cells were transfected with pSH-EFIRES-P-AtAFB2-weak-NLS and pCas9-sgAAVS1-1 (#129726, Addgene) and then selected in puromycin (5 μg/mL) after 4 days. Cells were then subjected to limiting cell dilution to obtain single colonies. These colonies were expanded, before genomic DNA was isolated and screened by PCR for the presence of the integrated AtAFB2-NLS transgene (see **Supplementary Table 1** for primer sequences).

A DNA plasmid used to target the AA7-3FLAG degron sequence to the C-terminus of *CTCF* (exon 12) was constructed in pUC19. To accomplish this, a 1,059 bp geneblock (Integrated DNA Technologies) representing a 500 bp left arm homologous to *CTCF* intron 11 and exon 12 and a 500 bp right homology arm of the *CTCF* 3’UTR, both flanking a multiple cloning site. The geneblock was flanked with *SalI/Acc65I* ends and silent mutations were also introduced to disrupt sgRNA complementarity and was cloned into pUC19 using *SalI/Acc65I* restriction endonucleases. A 345 bp geneblock (Integrated DNA Technologies) removing the *CTCF* stop codon and replacing it with a 3x ‘GGGGS’ flexible linker, the miniAA7 sequence and 3xFLAG tag, was cloned in using *AgeI/NotI* restriction endonucleases. A single-guide RNA (sgRNA), previously shown to efficiently direct Cas9 to a protospacer adjacent motif 9-11 bp after the *CTCF* stop codon (antisense strand), was cloned into pLKO.1-puro-U6-sgRNA-BfuAI-large stuffer (#52628, Addgene) which was modified to express an mCherry-2A-puro cassette from the human pgk promoter (see **Supplementary Table 1** for oligo sequences).

HEK293T-AtAFB2 cells were co-transfected with pLV-hUbc-Cas9-T2A-eGFP (#53190, Addgene), 52628-mCherry2Apuro-CTCF-sgRNA#4 and pUC19-CTCFex12-AA7-3xFLAG plasmids. After 3 days, cells were analysed by flow cytometry and enriched by FACS before single cells were isolated by limiting cell dilution. Individual clones were screened by dot blot for FLAG expression, before isolating genomic DNA and performing screening for the presence of the integrated AA7-3xFLAG template. Positive clones (e.g. 4.22 and 4.24) with all alleles edited were analysed by western blot to confirm FLAG expression and the absence of CTCF endogenous protein expression at ∼130 kDa.

### MTT assay

Cells were seeded in triplicate at appropriate densities into 96-well plate for each examined time point. Medium only control was included in triplicate in each plate. Cells were allowed to attach for at least 2 h. At each time point, 3-(4,5-dimethylthiazol-2-yl)-2,5-diphenyltetrazolium bromide (MTT, AG Scientific, USA) was added to each well and cells were incubated for a minimum of 4 h. After incubation, an equal volume of isopropanol and 2 M HCl was added to dissolve the resulting crystals. Absorbance was measured at 570 nm using a FLUOstar Omega microplate reader (BMG LabTech, Germany).

### Clonogenicity assay

Cells (300-1,000) were seeded in triplicate into 10-cm culture dishes with corresponding medium and incubated for 10-21 days to allow colony formation. Medium was changed after 7 days. At the experimental endpoint, samples were washed with PBS and then fixed in cold methanol at room temperature for 10 min. After fixation, methanol was aspirated and samples were allowed to air dry. Samples were stained with Giemsa Stain (Sigma-Aldrich, USA) for at least 30 min at room temperature and then washed 3 times in PBS. Colonies were counted and plates imaged using a Bio-Rad ChemiDoc imaging system.

### Statistical analysis

All experiments were conducted three times unless otherwise stated. Data is presented as mean ± SD with Welch’s t-test, one-way ANOVA with Dunnett’s multiple comparisons test or 2-way ANOVA with Šídák’s multiple comparisons test used for statistical analysis using GraphPad Prism where ns=not significant, *=*p*<0.05, **=*p*<0.01, ***=*p*<0.001 and ****=*p*<0.0001.

## Acknowledgements

The authors acknowledge funding to support the project from Tour de Cure, Tour de Rocks (C.B), National Health & Medical Research Council (NHMRC) Ideas Grant funding awarded to CGB (2037697, 2047345) and Cure the Future Foundation. Additional support was received from an NHMRC Project Grant (1128748) and an Investigator Grant (1177305) awarded to JEJR. The authors would also like to thank Sydney Cytometry for flow cytometry and imaging support.

## Contributions

Conceptualization, CGB and JEJR; data curation, SY and CGB; formal analysis, SY, SH, QS, CGB; methodology, SY, SH, RL, QS, MV, CM, MST and CGB; supervision, CGB; visualization, SY and CGB; draft, SY and CGB; writing – review and editing, SY, JEJR and CGB; funding acquisition, CGB and JEJR.

## Data accessibility

All analysed data have been provided within the paper.

## Supplementary Figure Legends

**Supplementary Figure 1.** Activation of ZNF185B expression after shRNA-mediated knockdown of *CTCF* in Ishikawa cells. **A)** Schematic of the dox-inducible pFH1tUTG-based vector used for shRNA-mediated knockdown of CTCF: long terminal repeat (LTR); H1 promoter with tetracycline operator (tetO); Ubiquitin-C promoter (UbC); tetracycline repressor (tetR); enhanced green fluorescent protein (eGFP); *Thosea asigna* virus 2A sequence (T2A). **B**) Flow cytometry analysis of KLE and Ishikawa cells following dox-inducible shRNA lentiviral transduction and fluorescent-activated cell sorting. The transduction efficiency (eGFP-positive cells) were quantified to determine the proportion of successfully transduced cells. **C**) Representative western blot of Ishikawa cell lysates after *CTCF* shRNA knockdown for 4 days which was probed with antibodies for CTCF, ZNF185 and APOE. GAPDH was used as a loading control, size markers (ladder, L) are indicated in kDa. **D**) Densitometric analysis of western blots from 3 independent experiments. Data represents the mean ± SD with statistical significance determined using 2-way ANOVA with Šídák’s multiple comparisons test with *=*p*<0.05, ***=*p*<0.001, ****=*p*<0.0001, ns=not significant.

**Supplementary Figure 2.** Validation of lentiviral overexpression of ZNF185 isoforms in Ishikawa cells. **A**) Flow cytometry analysis of Ishikawa cells following lentiviral transduction to examine transduction efficiency. Cells (eGFP positive) were quantified to determine the proportion of successfully transduced cells. **B**) Western blots of whole cell lysates from Ishikawa cells transduced with lentivectors expressing ZNF185A or ZNF185B were probed with antibodies against HA, universal ZNF185, ZNF185A and ZNF185B isoforms. GAPDH was used as a loading control and molecular weight markers are indicated in kDa. **C**) Immunofluorescence of Ishikawa cells transduced with lentivectors expressing ZNF185A or ZNF185B. Cells were stained with antibodies against ZNF185A isoform (red), ZNF185B isoform (red) or phalloidin (for F-actin, red), together with HA (magenta). Transduced cells were eGFP-positive (green), and nuclei were stained with DAPI (blue) in merged images. Non-transduced Ishikawa cells stained with antibodies against ZNF185 (red) and HA (magenta) were used as a negative control. Images were acquired with a 40× water immersion objective, scale bars = 25 µm.

**Supplementary Figure 3.** Effects of aberrant ZNF185 expression on endometrial cancer cell proliferation. **A**) Schematic of lentiviral vector used for shRNA-mediated knockdown of ZNF185: long terminal repeat (LTR); rev response element (RRE); U6 polIII promoter; central polypyrimidine tract (cPPT); phosphoglycerate kinase (pgk); enhanced green fluorescent protein (eGFP); picornaviral ribosomal skip sequence (2A); and puromycin resistance gene (puro) are indicated. The *ZNF185* mRNA sequence is depicted along with locations of shRNA complementarity. **B**) Representative clonogenicity plates of KLE cells transduced with *ZNF185* shRNAs after 21 days incubation and Ishikawa cells transduced with *ZNF185* shRNAs after 10 days incubation analysed in Figure 6G-H. **C**) MTT absorbance reading measured over 8 days in KLE cells transduced with pFUW-based lentivectors expressing empty vector (EV) or with ZNF185A or ZNF185B of n=3 independent experiments each performed in triplicate. Data represents the mean ± SD with statistical significance determined at the last time point using one-way ANOVA with Dunnett’s multiple comparisons test with *=*p*<0.05, **=*p*<0.01, ***=*p*<0.001, ****=*p*<0.0001, ns=not significant. **D**) Clonogenicity assay of KLE cells transduced with lentivectors expressing EV or with ZNF185A or ZNF185B after 21 days incubation. Colony number was normalised to EV. Data represents the mean ± SD with statistical significance determined using one-way ANOVA with Dunnett’s multiple comparisons test with ns=not significant. **E**) Representative clonogenicity plates of KLE cells transduced with lentivectors expressing empty vector (EV) or with ZNF185A or ZNF185B after 21 days incubation.

**Supplementary Table 1.**
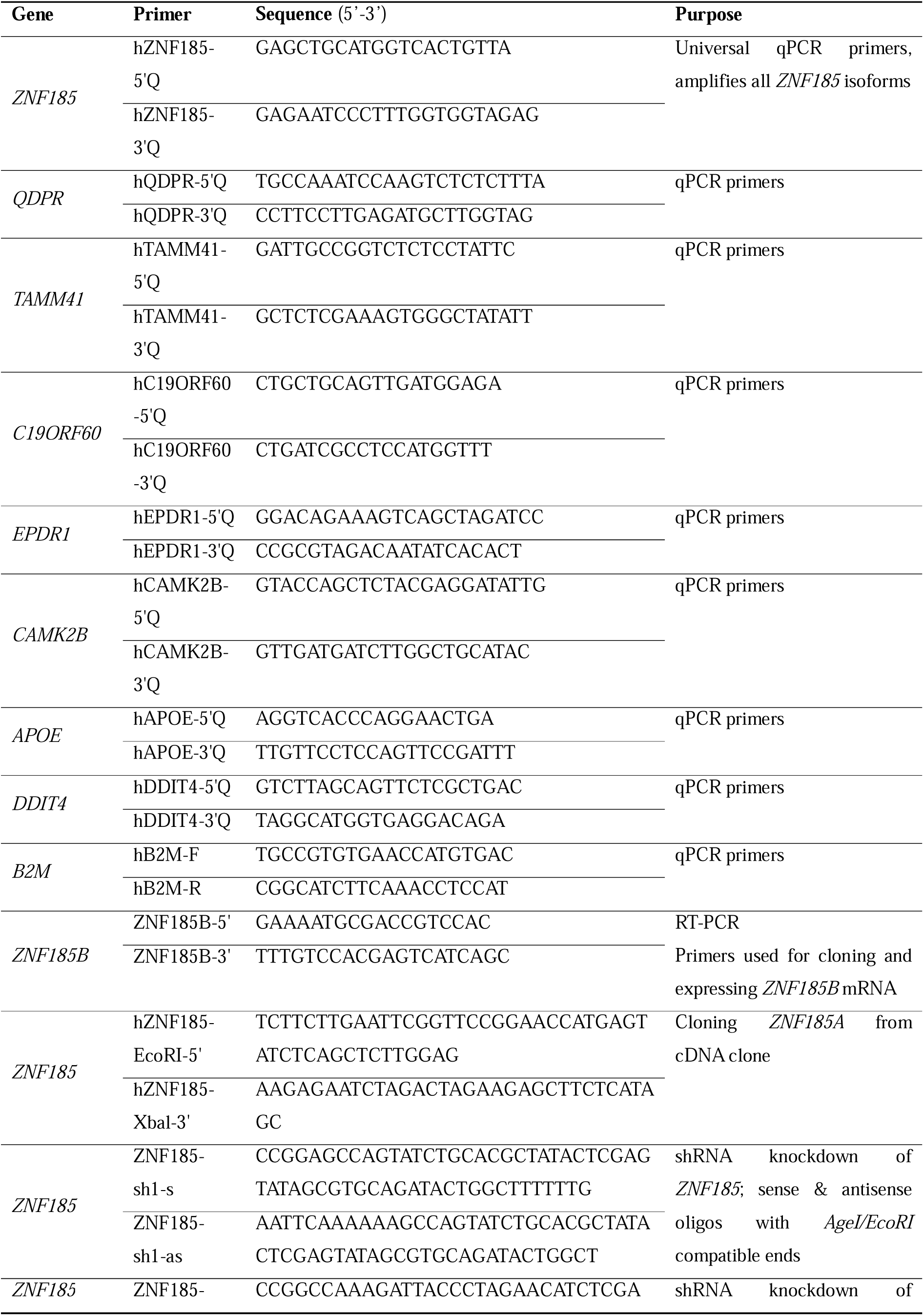
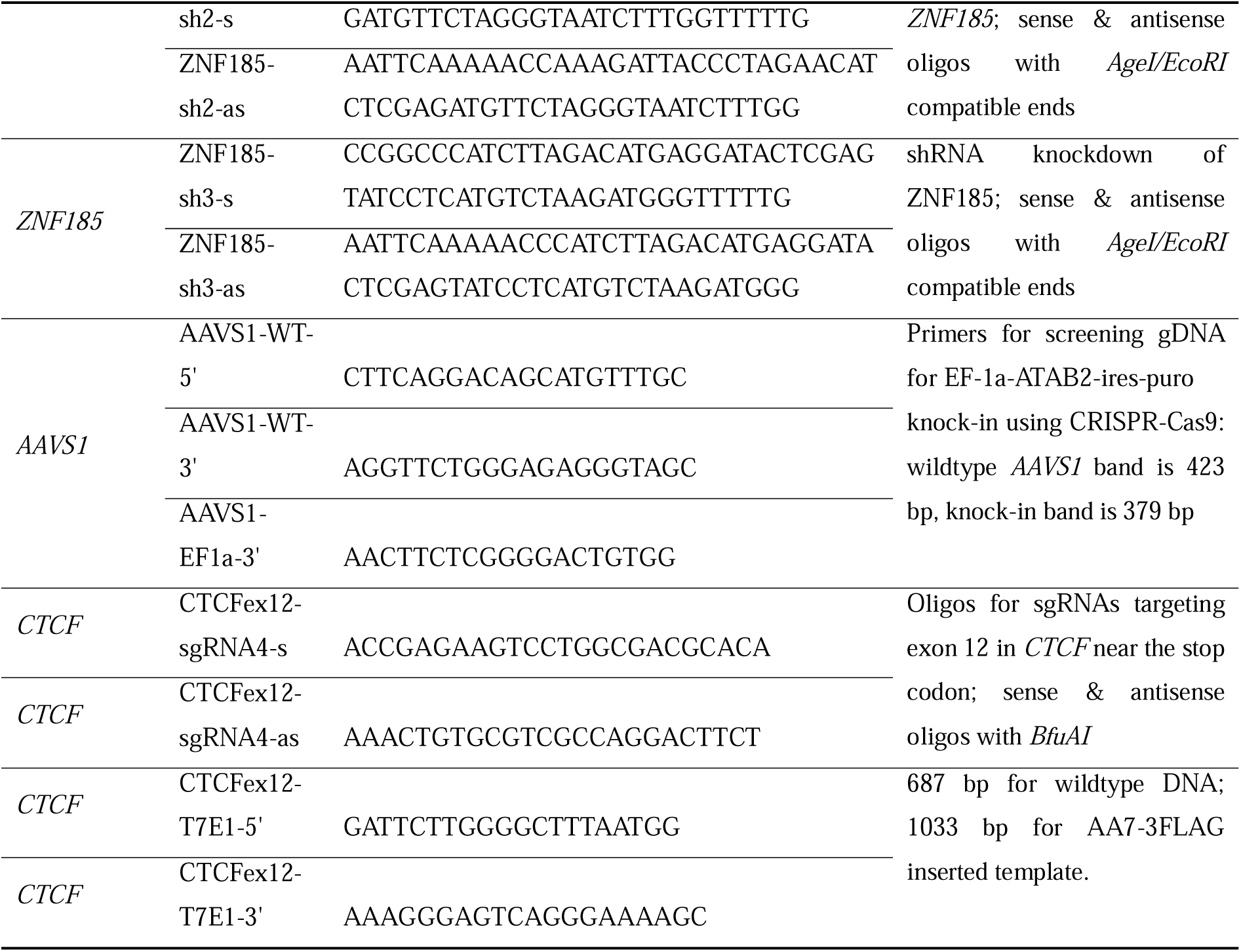
Oligonucleotides used in this study.

