## Supplementary figures and images for "ZNF185 expression is negatively regulated by CTCF and promotes endometrial cancer growth"

### Supplementary Figures

Supp. Figure 1

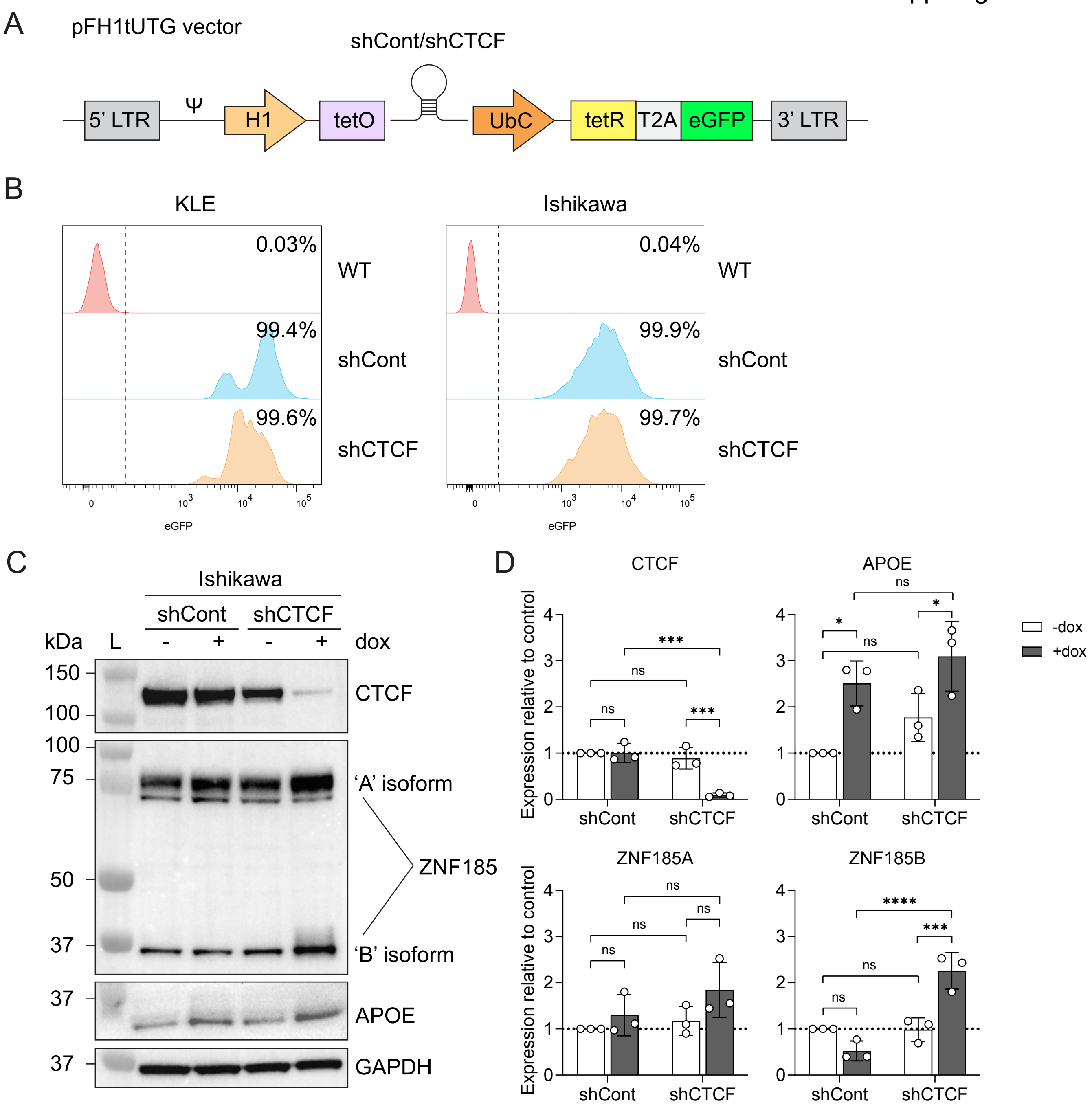

Supp. Figure 2

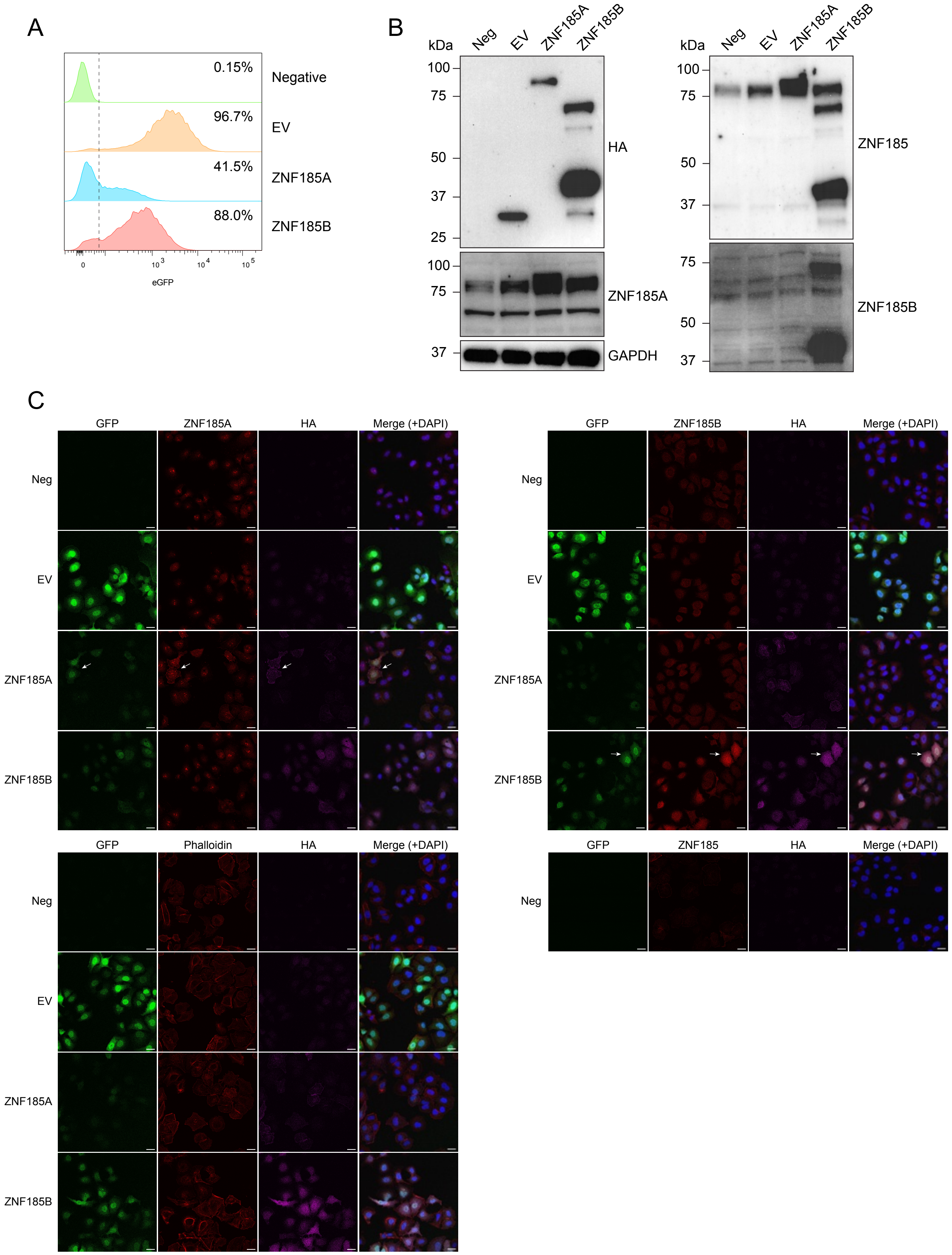

Supp. Figure 3

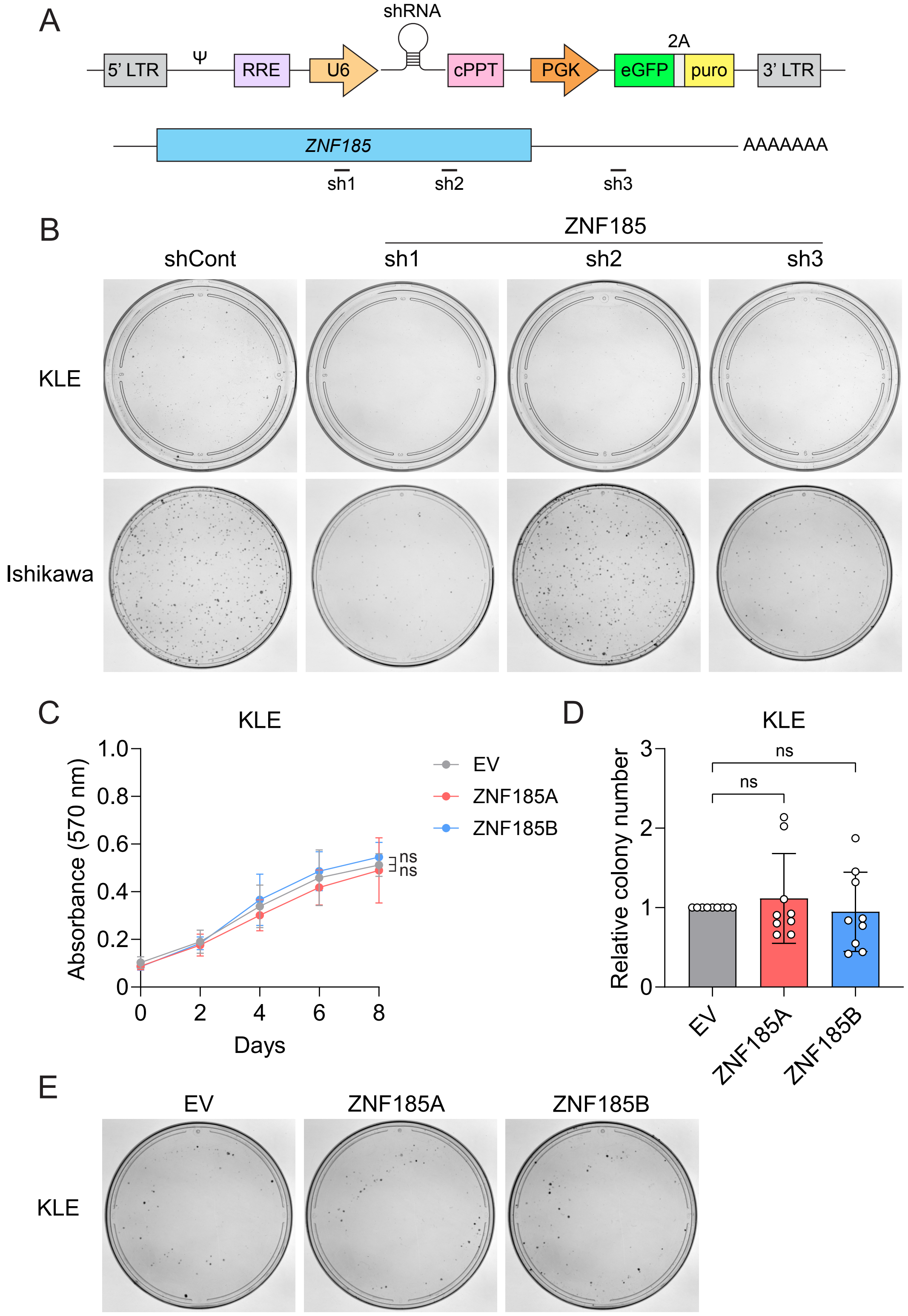
